# Protein-Protein Interaction Prediction is Achievable with Large Language Models

**DOI:** 10.1101/2023.06.07.544109

**Authors:** Logan Hallee, Jason P. Gleghorn

## Abstract

Predicting protein-protein interactions (PPIs) is vital for elucidating fundamental biology, designing peptide therapeutics, and for high-throughput protein annotation. This is particularly relevant in the current biotechnology landscape characterized by the proliferation of protein generative models, which necessitate a high-throughput and generalized PPI predictor for proteins regardless of conventional motifs or known biological functions. Our work addresses this need and provides strong evidence of the utility and reliability of protein language models (pLMs) in learning the PPI objective. We demonstrated that with the use of a sizable balanced dataset, pLMs achieve state-of-the-art performance metrics in PPI prediction on diverse proteins. To generate a dataset that allows for the approximation of these conditions, we implemented a novel synthetic data generation scheme to augment BIOGRID and Negatome datasets. The enhancement of these datasets was then used to fine-tune ProtBERT for PPI prediction to develop a model that we call SYNTERACT (SYNThetic data-driven protein-protein intERACtion Transformer). Our results are compelling, demonstrating 92% accuracy on validated positive and negative interacting pairs derived from 50 different organisms, all of which were excluded from the training phase. In addition to the high metrics, secondary analysis revealed that our synthetic negative data was able to successfully mimic actual negative samples, further reinforcing the integrity of synthetic data additions to PPI datasets. Another notable discovery was the ease in which previously existing PPI datasets could be predicted with simplistic features, calling into question if they can actually inform PPI prediction. We find that the subcellular compartment bias inherent to the compilation of these datasets is learnable with deep learning methods and demonstrate that our approach is not burdened by this disadvantage.

## 1 Introduction

Proteins are diverse macromolecules that manage the chemical reactions within biological systems. A protein’s structural and chemical properties dictate its function within a system, and their mutual interactions contribute to biochemistry. Dubbed protein-protein interactions (PPIs), these networks of interactions often pave the way to understanding biological processes. For example, DNA replication, transcription, translation, metabolism, and biological signaling[1, 2, 3, 4, 5, 6]. We define a PPI as physical contact that mediates chemical or conformational change, especially with non-generic function.

The gold standard for annotating PPIs is through “wet-lab” or *in vitro* validation, including mass spectrometry, yeast two-hybrid screening, microarrays, pull-down assays, and more[7, 1]. While these approaches deal with physical proteins, they are time intensive and expensive. With the remarkable advancements in generative artificial intelligence (AI) for biological applications, novel protein and peptide design offers high-throughput approaches to produce catalysts and therapeutics[8, 9]. Screening 100s of thousands or millions of proteins for a task is not feasible with *in vitro* methods. Instead, a highly accurate and robust *in silico* method is desirable to predict PPIs in a high-throughput manner.

The classic approach to predicting PPIs is to use existing structural data about individual proteins to make a prediction. Through Molecular Dynamics or graph-based neural networks, structural data often enables a robust prediction for interaction between two proteins[10, 11, 12, 13]. However, verified structural information is extremely sparse over the landscape of possible proteins due to the difficulty of obtaining crystal structures. Of course, deep learning (DL) approaches (AlphaFold2, RosseTTAFold, ESMFold, OmegaFold, EMBER2, etc.)[14, 15, 16, 17, 18] can provide high-throughput structural predictions with varying degrees of reliability. Unfortunately, designing a computational model from the output of another computational model compounds errors in an unsatisfactory way. Beyond structural availability, Molecular Dynamics simulations based on dense force fields are extremely computationally expensive, limiting the timescale of analysis. Multi-scale modeling seeks to combat this computational expense but is in the early stages[19, 20].

Fortunately, proteins can be represented more concisely than an atom-wise point cloud. Proteins comprise discrete standard units called amino acids, which can be organized and classified by their primary sequence: a string of letters where each unique amino acid has its own single-letter identity. Recent PPI predictors encode protein sequences with a numerical scheme and use DL techniques to make a binary prediction; interacts or not. These numerical schemes include known physicochemical features, one-hot encoding, amino acid composition, conjoint triad, auto covariance, and learned embeddings[1, 21, 22, 23]. However, the central problem with current PPI prediction is dataset availability; PPI data comes with a vast class imbalance[24]. Multi-validated interactors are published frequently, while validated non-interactors are hand-curated slowly. To combat this, researchers fit models to small, specialized datasets, assume random proteins from different subcellular compartments do not interact, or find random pairs that do not appear in interaction databases.

DL approaches can learn protein annotations such as subcellular compartment, solubility, or even phylogenetics[25, 26], and we hypothesize that many previous PPI models are actually partially learning the mechanism which researchers used to compile negative datasets and not a PPI objective. Thus, the aforementioned numerical encoding schemes result in high metric performance on these small or specialized datasets but could be learning the wrong objective. To develop a PPI predictor that can work on as diverse a variety of peptides as possible, we designed SYNTERACT (SYNThetic data-driven protein-protein intERACtion Transformer); a natural language processing (NLP) approach to PPI prediction that utilizes synthetic data generation to enable protein language model (pLM) fine-tuning with a balanced dataset. Using a pre-trained pLM, we leverage learned features to perform the PPI objective robustly, demonstrating 92% accuracy on validated positive and negative interacting pairs derived from 50 different organisms.

## 2 Methods

### 2.1 Datasets

#### 2.1.1 Training data

For positive interacting pairs, we used the multi-validated physical BIOGRID dataset version 4.4.213 (accessed August 30, 2022)[27]. Protein pairs (samples) qualify for the multi-validated physical dataset when proteins A and B are mapped as interactors by the same experimental methodology twice or using more than one methodology in separate publications. The list of possible validation experiments includes affinity capture mass spectrometry, biochemical activity (A modifying B), co-crystal structure, and co-purification, amongst others [27]. We mapped BIOGRID IDs to Swiss-Prot and Trembl accessions and sequences from UniProtKB (accessed October 14, 2022)[28]. We trimmed to sequence pairs resulting in a combined length of less than 1,000 amino acids for computational efficiency. The resultant positive interacting dataset was 179,018 sequence pairs.

Non-interacting (negative) pairs were downloaded from Negatome 2.0 (accessed March 18, 2023)[29], which were mapped and trimmed the same way as the positive interacting pairs. Negatome 2.0 is comprised of multiple groups, including protein pairs in the PDB that are members of a structural complex and do not interact directly, PDB structures filtered against IntAct, non-interacting PFAM domains found in the same complex, manually annotated literature (excluding high-throughput studies), manual curation vs. InAct, and manual curation of PFAM domain pairs[29]. This resulted in 3,958 non-interacting sequence pairs after preprocessing.

There were 160 species represented from our processed BIOGRID and Negatome datasets, with much of the protein data primarily comprised of homo sapiens and canonical animal models. The proteins ranged from 24 to 959 amino acids, and we preserved this diversity in our training data to produce as robust a predictor as possible.

#### 2.1.2 Synthetic negative data generation for balanced datasets

The glaring class imbalance in our dataset, with 179,018 positive and 3,958 negative interacting pairs, presents a significant challenge. With such disproportion, even a complex binary classifier would likely exhibit a misleadingly high accuracy of 99%+ by consistently predicting the majority class, i.e., positive interactors. This occurs due to the low probability of DL models identifying or learning from the significantly less represented negative interactors. To mitigate this issue, a balanced dataset with an equal distribution of positive and negative samples is typically preferred when training binary classifiers. Such a distribution neutralizes the model’s inclination to predict a class solely based on its prevalence. In this balanced scenario, a strategy of persistently predicting a single class, whether one or zero, would yield an accuracy of just 50%, equivalent to the outcome expected from random chance.

Three principal techniques can be employed to counteract imbalanced data in classification tasks: 1) Down-sampling the majority class, 2) Up-sampling the minority class through synthetic data generation, or 3) Implementing some combination of the two. In an effort to exploit as much available data as possible, we opted for the second strategy: to generate more than 170,000 synthetic negative samples for training our model.

Synthetic negative data generation was performed via two methods. Negative to negative: modifying a negative sample into a different negative sample, and positive to negative: changing a positive sample into a negative sample. Negative amino acid sequences were generated in both cases through BLOSUM-inspired substitutions. Evolutionary data informs the BLOSUM62 substitution matrix to understand the likelihood of mutations or substitutions between proteins. Positive scores are more likely to maintain common properties of a peptide after the substitution, whereas negative scores are more likely to be deleterious[30]. Of course, amino acid sequences are incredibly context-dependent, so these are not rigid rules but follow the trends of evolutionary lineage through the log-odds of aligned residues.

To generate a new pair of negative interactors from an existing negative sample, one protein was kept the same, and the other was mutated. A series of random positive scoring substitutions were applied. We chose positive mutations to reduce the likelihood of a spontaneous change in properties that enabled interaction with the other protein. To generate a new pair of negative interactors from an existing positive sample, one protein was kept the same, and the other was mutated. However, this time a series of random negative scoring substitutions were applied to introduce as many deleterious mutations as possible. The goal was to reduce the likelihood of maintaining the original interaction as much as possible. In both negative-to-negative and positive-to-negative cases, with some probability, we instead shuffled the adjusted sequence instead of mutating it. The assumption was that a shuffled protein is so jumbled that the likelihood of it preserving a meaningful interaction is very small. This pipeline is highlighted in **Figure 1**. Importantly, the starting methionine was never shuffled or mutated to preserve the common starting amino acid among most protein sequences.

**Figure 1:**
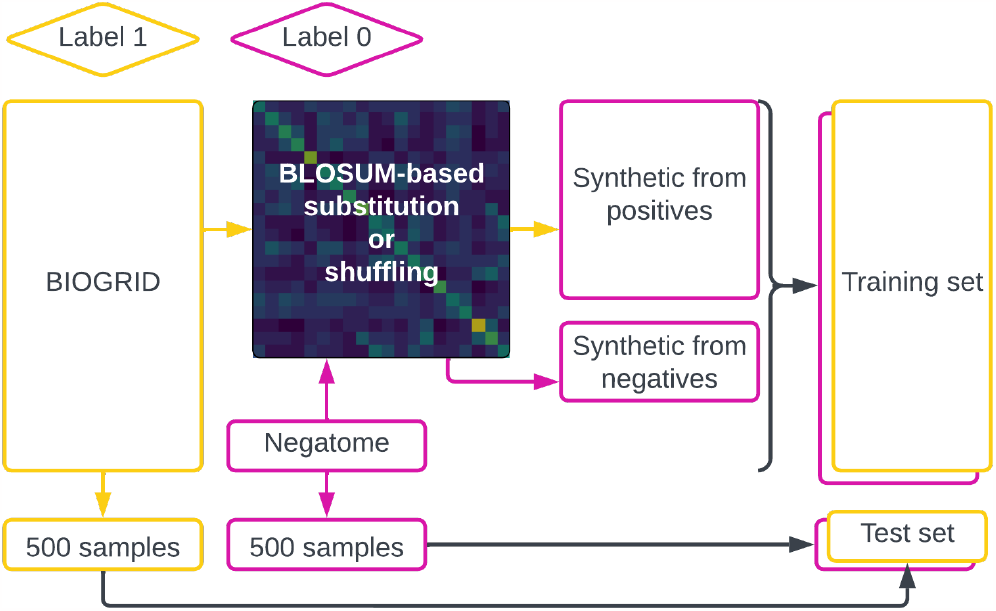
Data compilation from multi-validated BIOGRID and Negatome. Five hundred random samples were excluded from each dataset, and the remainder passed through our synthetic data generation pipeline. Most synthetic negatives were formed from modifying positive samples; however, both pathways resulted from specific sets of BLOSUM62-based substitutions or simple shuffling. Interacting samples were labeled one, and non-interacting samples were labeled zero.

#### 2.1.3 Test and validation data

A **test set** consisting of five hundred positive and five hundred negative samples were extracted randomly and held aside from synthetic data generation and model training. This test dataset was much smaller than desirable but was chosen to use as many Negatome samples as possible for training while having a balanced test set. There were 50 total species represented in the test set; however, most sequences were from model organisms. For evaluating the model performance during training, we used a **validation set**, a random 3% portion (10,000+ samples) of the training set that was not used to update the weights. We also utilized the newest BIOGRID release (**new BIOGRID set)** as of the time of manuscript preparation to further validate positive interaction prediction. BIOGRID 4.4.221 (accessed April 30, 2023) contained 4,534 positive interacting pairs that were not present in the BIOGRID version used for training.

We also downloaded and evaluated previously compiled datasets via the SDNN-PPI project GitHub[1]. These five datasets, Helicobacter pylori, Human Bacillus Anthracis, Human Yersinia pestis, Human, and Saccharomyces cerevisiae, were compiled with known positive interactors and random proteins from different subcellular compartments[1]. Throughout, we will refer to these datasets as **subcell datasets**.

Because multi-validated non-interactors are incredibly sparse and our test set only contained 500 negative examples, we sought to evaluate the non-interactor performance with additional methods. We hypothesized that the meaningful interaction rate of completely random proteins with each other is incredibly low. Random vertebrate mimetic proteins were generated to determine how well our model classifies non-interactors.

To generate random proteins, we sampled random amino acids at the frequency of non-membrane vertebrate proteins after a start methionine[31]. This approach was used to prevent bias from the amino acid frequencies between the proteins used to train the model and this evaluation set of protein pairs. They were of varying amino acid lengths (100 *< L <* 500), consistent with the vast majority of protein sizes. This procedure obviously removed common biologically relevant motifs that lead to common secondary structures, but artificial proteins are not bound by these restrictions. Due to the lack of common motifs, we expected this to be a straightforward task but necessary for our model. This group is referred to as **mimetic proteins**.

### 2.2 Protein language models

Advancements in understanding proteins based on primary sequence alone are attributed to the extensive use and continued understanding of transformer neural networks. Initially introduced in the NLP domain, transformers have proven highly effective in capturing complex patterns and dependencies within sequential data. Transformers learn effective numerical representations, called embeddings, through various mechanisms whereby unique sections of text, or tokens, are mapped to unique integers[32]. For pLMs the tokens are the amino acids, analogous to a “word” in typical NLP, and the entire protein sequence is a “sentence.” These integers serve as indices to access a predefined embedding matrix, essentially a lookup table. Each token is associated with a learned high-dimensional vector representation that captures both the syntactic and semantic information of the token.

A crucial component of transformer models is the multi-head self-attention mechanism, which captures long-range dependencies and relationships between tokens. Self-attention allows the model to weigh the importance of each token in the context of every other token; it tunes the embeddings in a context-dependent way. Mathematically, self-attention can be represented as follows:

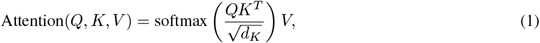

where *Q, K*, and *V* are the query, key, and value matrices, and *d*_*K*_ is the dimension of the *K*. For pLMs, the query, key, and value matrices are derived from the embeddings of the amino acids:

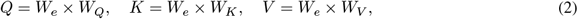

where *W*_*e*_ is the learned token embedding matrix (lookup table), and *W*_*Q*_, *W*_*K*_, and *W*_*V*_ are learned weights[32]. *K* and *V* are similar to how a traditional dictionary datatype works. The input amino acids are treated as the key, looking up values of these same amino acids and every combination thereof. *Q* is responsible for generating attention scores that determine the importance of each relationship between input amino acids. The final Attention output is a square matrix, *A*_*n×n*_, where *n* is the length of the input sequence and each index ranges from zero to one. Generally speaking, if *A*_*i,j*_ is large the *i*th amino acid is contextually important to the *j*th amino acid. Typically for pLMs, this implies a spatial or chemical relationship, such as the *i*th amino acid being close in 3D space to the *j*th amino acid of the actual protein[33]. *A*_*i,i*_ is always high.

For multi-head attention, the input sequence is transformed into multiple query, key, and value matrices, each corresponding to a different attention head with unique weights. Each attention head independently computes its version of *A*, which is concatenated into a final *n*×*n* matrix with an additional learned linear transformation[32]. Multi-head self-attention has been particularly important for pLMs, with higher head counts often showcasing higher performance at much higher compute costs[33].

In addition to attention, transformer models incorporate feed-forward layers, which introduce non-linear transformations to the encoded representations. These layers further enhance the model’s ability to capture complex patterns and dependencies within protein sequences[32].

Typically, a large corpus of text data is used to train transformers with masked language modeling (MLM)[34]. MLM is the process of randomly hiding portions of the text and then having the model “fill in the blanks,” repeating this many times. MLM offers a way to learn about massive corpora of amino acids in a semi-supervised manner that requires no human annotation, as the labels for training are the masked residues. The MLM task can be formatted as the following objective with a corpus of tokens *U* = *u*_1_, …, *u*_*n*_ maximizing the likelihood:

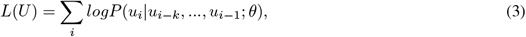

where *k* is the context window of masked tokens[34], and the conditional probability *P* is predicted by a transformer neural network with parameters *θ* (*W*_*e*_, *W*_*Q*_, *W*_*K*_, *W*_*V*_, etc.∈ *θ*). The output distribution of tokens is learned through the hidden or latent space outputs *H* = *h*_0_, …, *h*_*m*_:

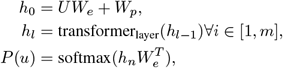

where *U* = (*u*_*−k*_, …, *u*_*−*1_) is the context vector of tokens, *m* is the number of transformer layers, and *W*_*p*_ is the position embedding matrix[34]. *L*(*U*) can be maximized during training by minimizing the cross-entropy between the model output and the original masked tokens. Once the weights have been “pre-trained” through MLM, optimizing weights for downstream tasks through supervised learning becomes much easier.

We used pre-trained weights from the ProtTrans project, which has conducted a massive amount of the MLM objective on various architectures[33]. pLM embeddings after MLM have been shown to describe protein physicochemical features. ProtTrans models have been fine-tuned to predict secondary structure, 3D structure, subcellular localization, binding residues, conservation effects of single amino acid variants, solubility, and more[33, 35, 36, 25, 37, 38]. Specifically, we used ProtBERT-BFD pre-trained weights to seed our model.

ProtBERT-BFD is the standard encoder-only BERT architecture but with 30 transformer layers, a hidden dimension of 1024, an intermediate dimension of 4096, and 16 attention heads[33]. It was trained on the Big Fantastic Database 100 (BFD), which contains over two billion protein sequences[14]. ProtBERT-BFD performs worse than the ProtT5-xl-uniref50 model on downstream tasks; however, it is much smaller. At 420 million parameters vs. ProtT5-xl at three billion, ProtBert-BFD is easier to fine-tune with limited data and compute. It is also easier for the deployment of downstream models.

To enable binary prediction with ProtBERT-BFD we tokenized the data in the format [*CLS*] Protein A [*SEP*] Protein B [*SEP*], where the [*SEP*] token allows the model to “separate” or distinguish protein sequences. The [*CLS*] token stands for classification and serves as an aggregate representation of the entire input [39]. We extract the [*CLS*] token embedding called the “pooled output,” and a feed-forward layer was added to project the final 1024-dimensional representation to a two-dimensional output. This was done by mapping the ProtBERT-BFD weights to the Huggingface BertForSequenceClassification module[40]. Softmax was applied to produce output logits to predict one (interacts) or zero (does not interact) with corresponding “confidences” for each prediction. **Figure 2** summarizes the final model architecture.

**Figure 2:**
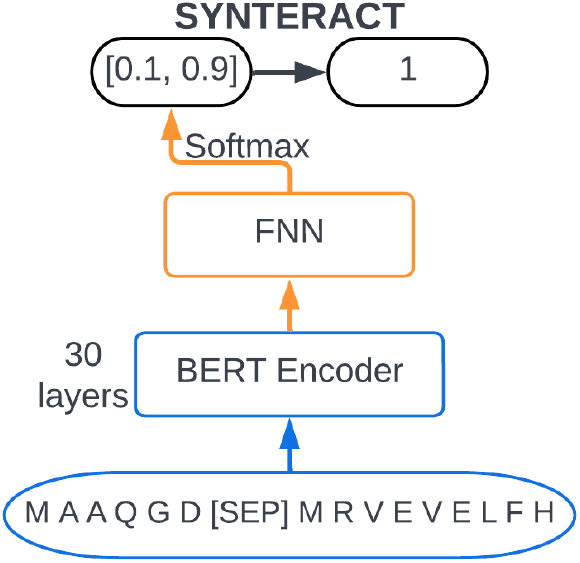
Summarized SYNTERACT architecture. SYNTERACT uses the 30 ProtBERT-BFD transformer encoder layers that pass to a feed-forward neural network comprised of a pooled representation and classification head. It was fine-tuned to take interacting proteins separated by [*SEP*] and output one or zero for interacting or non-interacting samples, respectively.

### 2.3 Training methodology

To train SYNTERACT, we kickstarted the fine-tuning of ProtBERT-BFD with the Huggingface autotrainer[40], allowing for the simultaneous training of 24 models. The PPI objective was difficult for the model to learn; many iterations presumably got stuck in local minima during gradient descent resulting in 50% or near 50% accuracy. However, using the AdamW optimizer’ stochastic nature, the best-performing model could get “lucky” with its weights, and perform with significantly higher metrics. We further trained the best-performing model for 20,000 steps with a global batch size of 70. Weights were saved when the model improved performance on the validation set.

For other training runs of SYNTERACT, for example, on subcell data or without Negatome samples mentioned below, we conducted a less expensive training to limit compute cost. They were similarly started with Huggingface autotrainer with 10 models and then trained roughly 30,000 steps with a batch size of 16 if necessary.

### 2.4 Support vector machines

When analyzing the possible objectives learned by DL PPI predictors, we utilized support vector machines (SVMs) as a proof-of-concept classifier. SVMs scale up the dimension of the feature data until a learned function can separate the labels. This function describes a hyper-plane, one dimension less than the feature data. Data that is well classified by SVMs is intuitively separable[26]. We used the standard hyperparameters in the sklearn python package for our experiments[41].

### 2.5 Protein feature extraction

Subcellular compartment and solubility of proteins were extracted through ProtT5-based subcellular compartment and solubility predictors; mapping each protein sequence to a two-dimensional feature space[25]. Therefore, for an input protein pair, there are a total of four features: one of nine unique subcellular compartment locations and solubility (membrane-bound or not) for both proteins. Each subcellular compartment and solubility class was assigned an integer.

Entire protein representations were extracted from the last hidden state of ProtBERT-BFD and SYNTERACT. This result was an array size *L* × 1024 where *L* was the length of both proteins for a sample together, including [*SEP*]. This array was the high-dimensional numerical representation of the proteins learned through MLM and fine-tuning. We took the average across the rows such that each protein pair input had a unique 1024-length vector representation.

## 3 Results

### 3.1 LLMs can learn PPI prediction

SYNTERACT performed with high metrics, including 92% accuracy on the test set from training-excluded multi-validated BIOGRID and Negatome datasets (**Table 1**). This diverse set of 50 different organisms highlights the robust nature of our primary-sequence-based predictions. When new BIOGRID samples were input into SYNTERACT, the model produced similar metrics (96% accuracy) demonstrating a high predictive power on samples that were not annotated at the time of training. Similarly, when SYNTERACT evaluated the entire Negatome, the model achieved 96% accuracy. Evaluating on the entire Negatome included examples trained on and excluded, thus, because this performance was similar to that of the new BIOGRID samples we conclude that the model is not overtrained. We also evaluated on random vertebrate mimetic proteins that we generated and SYNTERACT achieved 100% accuracy. Complete confusion matrix data is reported in **Figure 3**.

**Table 1:**
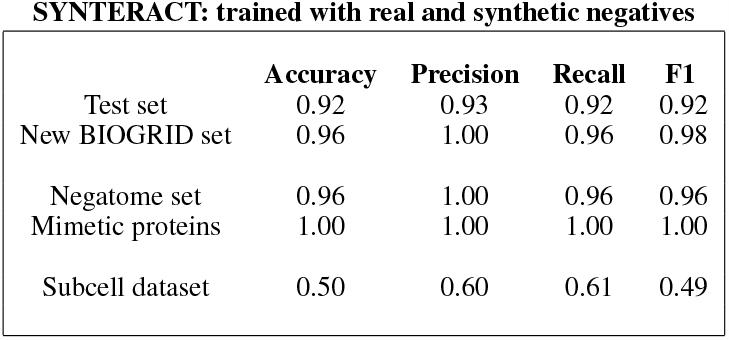
SYNTERACT model trained on BIOGRID, Negatome, and synthetic negatives. Performance metrics of accuracy, precision, recall, and macro-average F1 score of SYNTERACT evaluated on other datasets. A high-scoring metric is close to 1.

**Figure 3:**
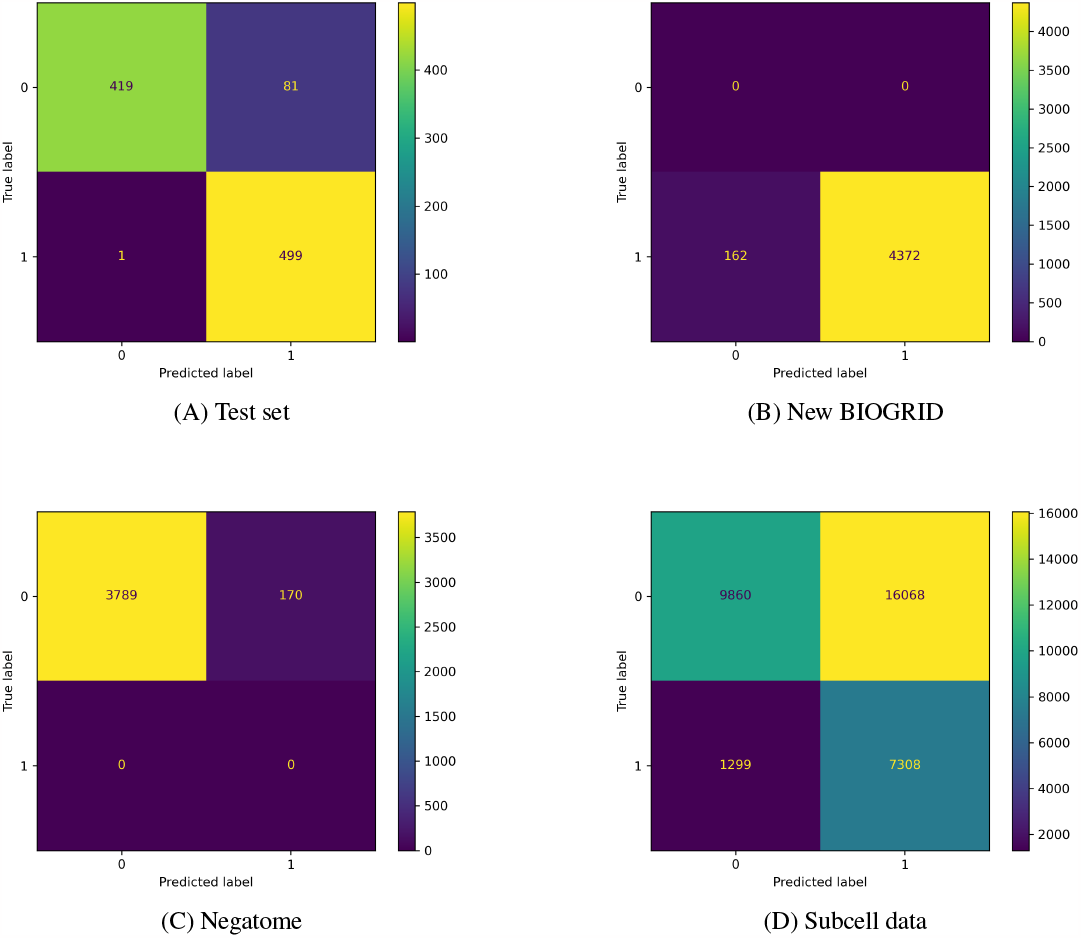
Confusion matrices of results for SYNTERACT trained on our dataset with real and synthetic negatives. Correct results are indicated where the true and predicted labels match, along the left-right diagonal. (A) Our test dataset. (B) New BIOGRID examples that did not exist during dataset compilation. (C) Full preprocessed Negatome, including training and test data. (D) Human, human helicobacter pylori, human bacillus anthracis, human yersinia pestis, and saccharomyces cerevisiae subcell datasets combined.

### 3.2 LLMs can learn negative generation schemes

We evaluated performance metrics on the subcell data set consisting of previously compiled human, human helicobacter pylori, human bacillus anthracis, human yersinia pestis, and saccharomyces cerevisiae with known positive interactors and random proteins from different subcellular compartments as negative interactors. On positive samples in this dataset, the model was 85% accurate. On negative samples, SYNTERACT was only 38% accurate, classifying many negatives from subcellular compartment sampling as interactors (**Figure 3**). Because SYNTERACT performed significantly worse on these datasets compared to validated interactions, especially on negatives, we were suspicious of how previous negatives have been compiled.

Current approaches to generating negative interactors include assembling non-interacting protein pairs from different subcellular compartments. We find the practice of choosing random proteins from different subcellular compartments a suboptimal approach to generating non-interacting samples; for the simple reason that in the cell, many proteins from separate compartments do colocalize and interact and even if they do not conventionally interact *in situ*, they could *in vitro*. Additionally, *we suspect that artificial patterns that arise from randomly pooling proteins from different subcellular compartments are a learnable objective for a machine-learning model*.

To test this hypothesis, we extracted subcellular compartment and solubility features from the human subcell data. We used the four features for each pair to predict their zero and one interaction labels with an SVM. There were only 324 unique combinations of those classes, so there were bound to be duplicates between a training and test set. As such, we only evaluated on the training data. We chose this approach as a proof-of-concept to illustrate how simple patterns can emerge from how researchers curate PPI datasets. This process was also repeated on a subset of our data for comparison.

Of course, how we synthetically generated data could also lead to obscure biases and skewed model metrics that do not correctly represent a PPI objective. To determine if Negatome samples could be properly classified with only knowledge of our synthetic non-interacting samples, we trained a model from scratch with no Negatome or with Negatome-derived protein pairs. We fit an SVM classifier on the embeddings of our training set to determine if such a classifier could classify Negatome data with only knowledge of real positives and synthetic negatives.

To extract embeddings for our SVM classifier, we ran 25,000 real positives and 25,000 synthetic negatives. We fitted an SVM on these embeddings to classify our labels and evaluated on our test set constructed from BIOGRID and Negatome samples (**Figure 4A**). This was repeated for an untrained SYNTERACT (the same weights as ProtBERT-BFD) and our trained SYNTERACT. We also trained SYNTERACT from scratch on the subcell data and validated on test subcell data (**Figure 4B**), trained SYNTERACT on the subcell data and validated on our test data (**Figure 4C**), and evaluated our trained model on the subcell data (**Figure 4D**). This comparison of metrics allowed us to discern whether the objectives learned from the subcellular data align with those learned from our own data. They provide insight into whether both sets of objectives are tractable using SYNTERACT and if they are distinctly different from one another.

**Figure 4:**
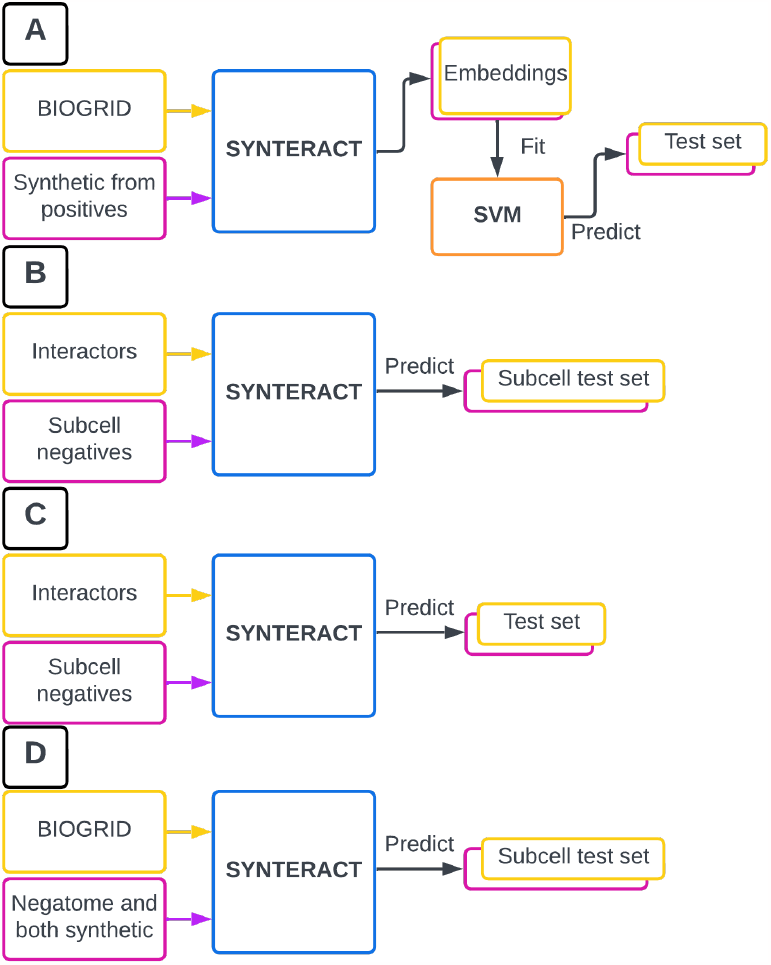
Scheme for comparing objectives. A. Validation that a dataset of synthetic negative and true positive embeddings can be used to predict real positives and real negatives. B. Determining if SYNTERACT trained on subcell data can perform well on a test subset. C. Determining if SYNTERACT trained on subcell data can perform well on our test data. D. Determining if SYNTERACT trained on our data can perform well on subcell data.

Model performance of SYNTERACT, when retrained on human subcell data, yields 96% accuracy on a test set of 10% of the samples excluded from training (**Table 2**). However, the same model performs poorly on our multi-validated test set with only 54% accuracy, marginally better than random chance. When repeated with all five subcell datasets combined, the results follow a similar trend but lead to a more robust model. SYNTERACT trained on all five subcell datasets performs with 64% accuracy on our test set and 87% on its own test set (10% of the total samples excluded from training) (**Table 2**). This suggests that the objectives learned from the varying types of negative samples are distinctly different.

**Table 2:** SYNTERACT trained on subcell data sets. Performance metrics of accuracy, precision, recall, and macro-average F1 score of SYNTERACT evaluated on different test datasets.

|  | **Accuracy** | **Precision** | **Recall** | **F1** |
| --- | --- | --- | --- | --- |
| Human subcell test set (trained on human subcell data) | 0.96 | 0.98 | 0.93 | 0.95 |
| Our test set (trained on human subcell data) | 0.54 | 0.53 | 0.71 | 0.61 |
| All subcell test set (trained on all subcell data) | 0.87 | 0.83 | 0.82 | 0.83 |
| Our test set (trained on all subcell data) | 0.64 | 0.64 | 0.64 | 0.64 |

When the subcellular compartment and solubility-based prediction was conducted on our test set, the SVM classifier resulted in 50% accuracy, poorly classifying our zeros and ones (**Table 3**). However, the human subcellular data was classified considerably better with a 65% accuracy. More importantly, the SVM model trained on human subcellular data correctly classified 95% of the zero labels, indicating that this sampling strategy might promote a learnable pattern.

**Table 3:** SVM model results after being trained on small four-feature versions of our test dataset and the human subcell dataset. The membrane and solubility features do not classify our data well but appear to classify subcell data partially.

|  | **Accuracy** | **Precision** | **Recall** | **F1** |
| --- | --- | --- | --- | --- |
| Our data | 0.50 | 0.25 | 0.50 | 0.33 |
| Subcell data | 0.65 | 0.72 | 0.63 | 0.60 |

### 3.3 Our synthetic negatives mimic non-interactors

Training SYNTERACT from scratch with no Negatome or Negatome-derived samples did not lead to the model learning a strategy that performed significantly over 50% on the test data (**Table 4**). This suggests that some number of actual negative samples are necessary to teach pLMs the PPI objective.

**Table 4:** SVM model results trained on embeddings from SYNTERACT. With the original ProtBERT-BFD weights, the synthetic negative embeddings were insufficient to differentiate between true positives and true negatives. After training on our dataset, synthetic negative embeddings provide enough variance to classify true negatives partially.

|  | **Accuracy** | **Precision** | **Recall** | **F1** |
| --- | --- | --- | --- | --- |
| From scratch | 0.50 | 0.50 | 0.50 | 0.39 |
| From trained embeddings | 0.73 | 0.81 | 0.73 | 0.71 |

Unsurprisingly, ProtBERT-BFD embeddings of 50,000 BIOGRID and synthetic BIOGRID-derived negatives with no PPI training resulted in low classification metrics on our test dataset with 50% accuracy. However, when using the embeddings from the trained SYNTERACT this same data was used to fit an SVM with a significantly higher accuracy (73%), misclassifying six positive pairs and 262 negative pairs. Importantly, this SVM correctly classified 238 real Negatome pairs without ever “seeing” real non-interacting pairs. This implies that our synthetic data generation at least partially represents true non-interactors.

## 4 Discussion

### Our experiments demonstrate that with a large balanced dataset, pLMs can successfully learn the objective of PPI prediction with high metrics robustly

Because non-interactors are largely unavailable compared with positive interactors, we synthetically generated negative samples to approximate these conditions for success. Whereas embeddings from real interactors and our synthetic non-interactors can partially predict Negatome samples, our model falls short of high-scoring performance without the addition of Negatome examples. However, we conclude that the model is not overtrained, performing similarly on samples that were included (Negatome dataset) and excluded (new BIOGRID dataset) from the training process.

Importantly, our dataset did not undergo homology-based trimming. Although this would be a preferable way to prevent the memorization of specific samples or motifs, our decision was driven by the resultant severe depletion of usable Negatome samples and a significant reduction in BIOGRID samples. This issue is further amplified when extended from trimming based on UniProt ID to some percentage of the sequence identity. Due to the uneven focus of biological research on specific proteins, there were unavoidable repeats of single proteins across training and test sets. However, there was no single protein pair (A + B) matching between the train and the test set. Luckily, many proteins appear in both Negatome and BIOGRID samples. Since we derived synthetic negatives from both sets, finding a protein exclusive to positive or negative labeling was rare - i.e., protein A was part of an interacting pair sometimes and a non-interacting pair at other times. This implies that SYNTERACT likely avoided memorization of proteins as strictly interacting or non-interacting, as they were inconsistently labeled across the training set. A similar inconsistency was observed in the test set, where a small subset of proteins appeared in both the interacting and non-interacting samples.

We suspect that the practice of generating negative samples by random selection from different subcellular compartments caused DL-based PPI predictors to learn an incorrect objective. We support this claim by showing that subcellular compartments and solubility alone have substantial predictive power on previously established datasets without any information on the sequence in question, particularly in negative samples. Our dataset does not have this potential objective because our labels are poorly predictable from the subcellular compartment and solubility alone. Additionally, SYNTERACT trained on subcell data can predict test subcell data but poorly predicts multi-validated positive and negative samples from BIOGRID and Negatome.

Of course, our final model is likely not without bias either. When evaluated on randomly generated vertebrate mimetic proteins, constructed similarly to our synthetic data non-interactors, our model was 100% accurate. The high level of accuracy prompts careful interpretation of the underlying causes. We assume that proteins rarely interact randomly; usually under specific physiological contexts guided by cellular machinery. The 10,000 randomly generated protein pairs, therefore, may not reflect the variety and complexity of interactions found within biological systems. Furthermore, the random sequences are unlikely to contain naturally common motifs that are crucial for meaningful interaction. Thus, the interaction rate of random proteins may truly be less than one in 10,000, but our model could also be biased towards random sequences due to the introduction of shuffled sequences in training.

Unfortunately, our false-positive predictions were high when evaluated using subcell data. We suspect that many of the negative samples from subcell data do, in fact, interact, but our model still predicted interactors on these datasets at a higher-than-expected rate. Our goal was to determine the PPI objective that predicts physical contact that mediates chemical or conformational changes, especially with *non-generic function*. Because of its performance on subcell data, we suspect that SYNTERACT has only learned some notion of physical contact that mediates chemical or conformational changes with or without a *biologically relevant function*. We still greatly value this more generalized objective because artificially generated proteins do not have a naturally occurring biological function.

Our high-performance metrics on multi-validated positive and negative interactors show that pLMs are a valuable tool for difficult protein annotation problems. Their broad capability in amino acid space and sequence size opens the possibility of predicting interactions of small peptide therapeutics, protein domains, or even entire biological pathways. We hope our model lays the foundation for primary sequence pLM-based high-throughput annotation of proteins and peptides, including artificial proteins. We see immense value in the continued curation of non-interactors for further informing pLMs to predict interactions.

## 5 Funding and Acknowledgments

This work was partly supported by the University of Delaware Graduate College through the Unidel Distinguished Graduate Scholar Award (L.H.) and the University of Delaware Artificial Intelligence Center of Excellence (AICoE). Any opinions, findings, and conclusions or recommendations expressed in this material are those of the authors. We would like to acknowledge the valuable feedback and critical review of this work from Yasaman Moghadamnia, Krithika Umesh, and Yuanjun Shen Ph.D., along with Aaron Oster for his help with the synthetic generation code. Figures were generated with Lucidchart, www.lucidchart.com.

